# Flame-forged divergence? Ancient human fires and the evolution of diurnal and nocturnal lineages in Mediterranean geckos

**DOI:** 10.1101/2024.03.05.583480

**Authors:** Domenico Fulgione, Danilo Russo, Eleonora Rivieccio, Valeria Maselli, Bice Avallone, Alessandro Mondanaro, Giorgio Giurato, Maria Buglione

**Affiliations:** Department of Biology, University of Naples Federico II, Naples, Italy, 80126; Department of Agriculture, University of Naples Federico II, Portici (Na), Italy, 80055; Department of Humanities Studies, University of Naples Federico II, Naples, Italy, 80138; Department of Earth Sciences, University of Florence, 50121; 5Department of Medicine, Surgery and Dentistry Medicine, University of Salerno, Salerno, Italy, 84081

**Author notes:** Correspondence to Danilo Russo Department of Agriculture, University of Naples Federico II, Reggia di Portici, Piazza Carlo di Borbone, 1 - 80055 - Portici (NA).

## Abstract

Human influence has historically exerted a major driving in creating novel ecological niches. Although the controlled use of fire by ancient humans probably played a significant role by attracting positive phototactic prey and favour foraging by insectivorous vertebrates, no study has ever explored this possibility. Using a multidisciplinary approach, we explore whether human-controlled fire has historically affected the temporal niche partitioning in two lineages of Moorish geckos *Tarentola mauritanica*, a diurnal- dark form and a nocturnal-pale form. We showed that the nocturnal-pale variant possesses lower skin melanin for fewer and smaller melanosomes and experiences lower α-MSH plasmatic levels than its diurnal-dark counterpart. Additionally, the analysis of the full mitochondrial genome established that our pale-nocturnal lineage emerged around 40,000 years ago, i.e., 20,000 years before the dark-diurnal lineage. Both variants arose when modern humans expanded into Europe, coinciding with the widespread use of fire, which likely facilitated the availability of arthropod prey for pale geckos, opening a previously untapped niche. Our modelling exercises corroborated a highly probable human-geckos coexistence during the emergence of these lineages. Furthermore, we experimentally demonstrated that fire near a rock surface significantly increases the abundance of arthropod prey, attracting preferred prey for the pale variant during nocturnal foraging. We suggest that ancient fires likely provided a novel foraging niche for pale geckos, yet it remains unclear whether the two lineages originated from diurnal or nocturnal ancestors. We present two alternative interpretations: the “out of the dark” scenario proposes a nocturnal ancestor leading to a diurnal descendant, while the “into the dark” scenario suggests a dark-diurnal ancestor evolving into a pale-nocturnal lineage. Phenotypic plasticity emerges as a critical factor in both scenarios, facilitating adaptation to new environments. We underscore the ongoing impact of artificial lighting on nocturnal behaviour, offering parallels to the potential origins of the gecko’s nocturnal lineage.

## Introduction

Humans are the only animal species capable of changing the planet on a global scale [1]. Our presence has largely affected the fate of wildlife throughout the history of humankind, promoting vast changes in community composition and geographic ranges [2]. Humans have also caused numerous extinction events [1,3], filtering out many species from human-altered landscapes where more opportunistic species persist or thrive [4,5], and not infrequently exerting selective pressures that have led to new phyletic lineages and speciation [6].

Artificial lighting at night (hereafter, ALAN) is one of the most pervasive forms of environmental alteration caused by humans. Numerous organisms adjust their behaviour and life cycles according to the availability of light across various time frames, ranging from daily to seasonal cycles. By introducing changes to the natural cycle of light and darkness, ALAN significantly impacts several crucial aspects of animal existence [7,8], especially affecting nocturnal species, whose sensitivity is evident as they either actively avoid light, as seen in most bat species [9,10], or are attracted to it, as in numerous nocturnal insects showing positive phototaxis (e.g., [11].

While the large-scale, overwhelming impact of ALAN on animals is well-known (e.g., [12], the effects of the first form of human-generated lighting, fire, are practically unknown.

Humans have been using fire for hundreds of thousands of years: the intentional use of fire traces back to the Middle Pleistocene of the Levant [13,14] but understandably spread as a common practice just afterwards [15]. Whereas the earliest fire usage may suggest cooking and anti-predatory activities, the more widespread use of fire to repel predators and allow extra-social interactions may have led to a “daylight extension”, in which humans have developed an activity peak during late evening hours, an unusual condition compared to the mammalian standard, including other primates [16]. Such a change in the temporal niche may have been linked to the development of language during the latest Pleistocene [17].

Not only may fire change the microclimate, but it also brings about light, including at night, when numerous phototactic insects may get lured and potentially fall victim to opportunistic predators exploiting the new environmental conditions and their associated food bonanza. However, no study has explored the possibility that fire has historically generated a previously untapped foraging niche suitable for insectivorous vertebrates. Although there is no way of retrospectively investigating the soundness of this hypothesis, several clues may be gathered at least to test the potential influence of ancient human- governed fires on animal ecology and evolution. To reach this goal, we selected the Moorish gecko (*Tarentola mauritanica*) as a case study. This small saurian is widespread in North Africa and Europe and has the widest range in the Mediterranean Basin of all gecko species [18]. Besides, because of the close relationship with humans, the species has been accidentally introduced to the Balearics (Spain), Madeira (Portugal), some Balkan islands, Crete, and South America [19,20]. European populations of Moorish geckos diverged during the early Pliocene, around 4.14 Mya. The colonization of the Iberian Peninsula by Moorish geckos from North Africa could have occurred during the Messinian Salinity Crisis or after the opening of the Strait of Gibraltar during the early Pliocene [21].

*Tarentola mauritanica* is active during both diurnal and nocturnal hours, occupying a variety of ecological niches, such as tree trunks, houses, and stone walls [22–24]. This species represents an interesting example of plastic variation, highlighted by its ability to adapt its skin colour according to the substrate [25]. Two distinct and sympatric populations can be easily distinguished [22] (Figure 1): the “dark diurnal gecko,” which mainly lives on trees, and the “pale nocturnal gecko,” found in human settlements on walls, especially near artificial lights, to increase the chances of prey capture [22,26,27]. The different colourations (Figure 1) are adaptations to improve camouflage during the day (favoured by the dark form) or at night (favoured by the pale form), as established experimentally [22].

**Figure 1.**
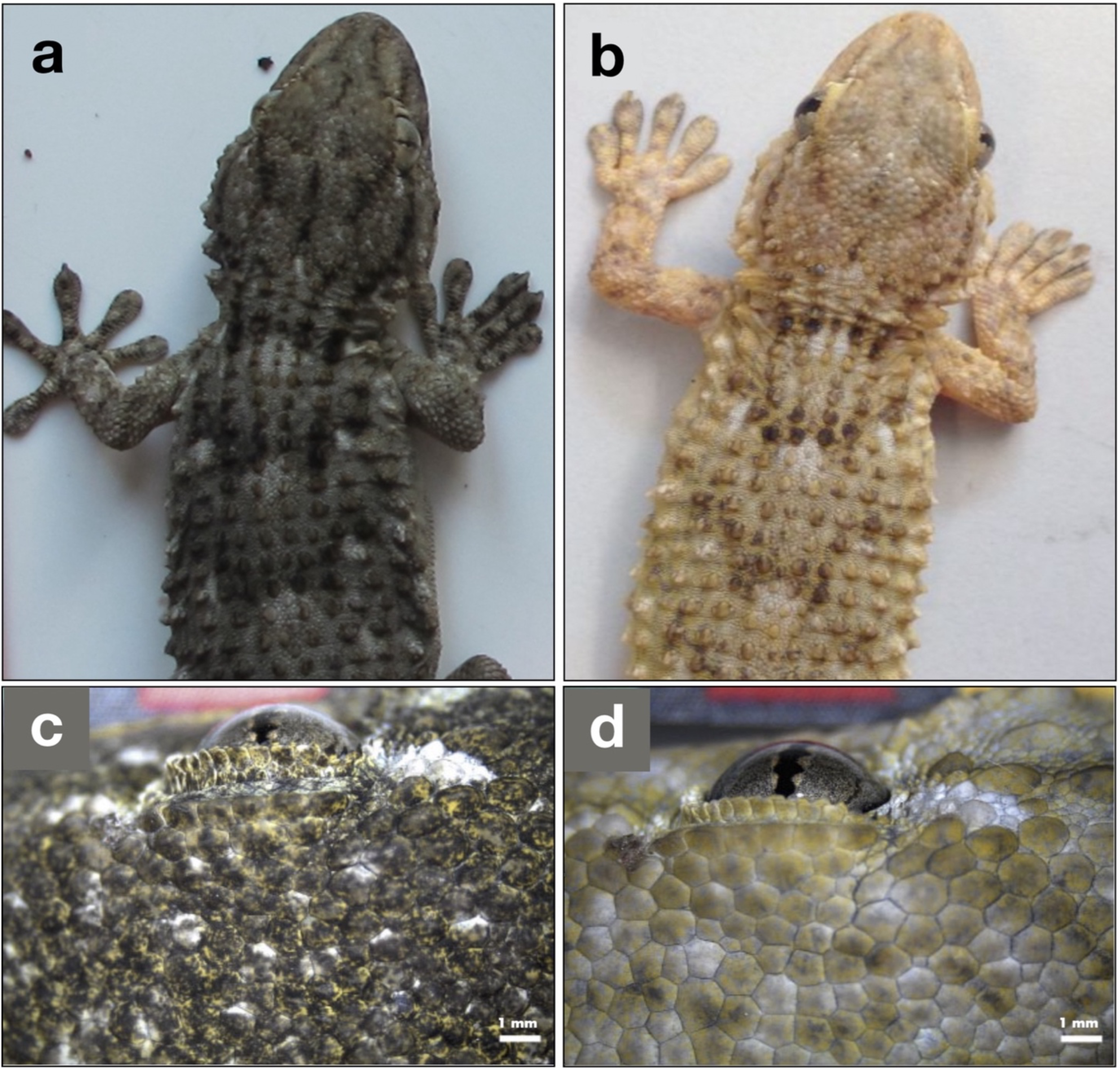
Dark-diurnal (a) and pale-nocturnal (b) Moorish geckos (*Tarentola mauritanica*). The two lineages occupy distinct ecological niches: the diurnal form inhabits trees, while the nocturnal variant prefers building walls during the night. Insets (c) and (d) show the plasticity of a dark-diurnal gecko changing skin colour on different substrates (see [25] for more details).

We propose that as modern humans spread across Europe and especially the Mediterranean, human-controlled fire played a significant role, introducing a new foraging opportunity for Moorish geckos. This opportunity arose from positive phototactic arthropods being attracted to the light provided by fire. It is currently unknown whether nocturnal Moorish geckos are the ancestral form from which diurnal geckos have evolved, or if diurnal geckos represent the ancient diurnal form that has subsequently adapted to the nocturnal niche. We argue that both evolutionary paths are conceivable and may be associated with the spread of human-generated fire. This association can be illustrated by the following scenarios:

*”Out of the dark” scenario*: The ancestral form was nocturnal and largely exploited human-generated fire. However, Moorish geckos are highly territorial and aggressive [28,29], so only some dominant individuals controlled the food bonanza offered by fire, whereas less competitive individuals were excluded from such profitable foraging sites and confined to peripheral sites whose food availability was impoverished by the “vacuum effect” exerted by the fire. The more phenotypically plastic geckos exploited their capacity to adapt their colour to the diurnal conditions and mitigate predation in daylight, so they conquered a novel diurnal niche, leaving the less plastic, dominant individuals associated with humans and their fires.

*”Into the dark” scenario*: The strong plasticity in skin colour shown by this species has favoured the evolution of a night-adapted lineage and led to the split between dark and pale geckos. The direct ancestors of the nocturnal form may have consisted of individuals showing an especially pale colour that protected them from nocturnal predators [22] while allowing them to select human fires set in caves or near rocks as successful foraging habitats. By attracting positively phototactic insect prey, illuminated cave walls may have offered a new foraging niche that has been successfully exploited by the geckos, leading to the evolution of a paler, nocturnal, and synanthropic lineage that has ever since accompanied our species and the creation of urban settlements up to modern times.

*To explore these evolutionary scenarios*, we formulated the following hypotheses and predictions:

1. Although to some extent the two forms, diurnal and nocturnal respectively, both retain the ability to vary the level of darkness/lightness [22], nocturnal geckos are characterised by a more marked depigmentation resulting from their high adaptation to the nocturnal niche. We, therefore, predict that a) the skin of the two forms will differ in the size and numerosity of melanophores (the cell types responsible for skin colour) as well as skin reflectance and that b) the blood levels of α- MSH, the hormone that stimulates melanin production in the dermis, will be significantly lower in nocturnal pale geckos than in diurnal, dark geckos, overall revealing distinct structural features between the lineages that are adapted to the two temporal niches.
2. Modern humans emerged in Europe, particularly within the Mediterranean region, during the Late Pleistocene, as supported by evidence from Italy and Bulgaria dating approximately 45-43 thousand years ago [30–32], and from France, dating back to 54,000 years B.P. [33]. The advent of modern humans brought about the widespread use of fire, potentially creating a unique foraging niche for a specialised gecko lineage. It is plausible that Moorish geckos and humans encountered each other in the Mediterranean region during this period, assuming they inhabited the same areas. Consequently, we hypothesise that the divergence of the two lineages occurred during the period of sympatry between humans and Moorish geckos. We anticipate that this timeframe will align with the estimated splitting time derived from molecular dating in phylogenetic analysis, with potential sympatry confirmed by species distribution models projected to that time. Furthermore, the “into the dark” scenario would imply an older origin of the diurnal lineage compared to the nocturnal lineage, whereas the opposite would hold according to the “out of the dark” scenario.
3. Fire may significantly increase prey availability when lit near rock surfaces by attracting insects that are eaten by the nocturnal form of the gecko. Preliminary work has established a clear diet difference between diurnal and nocturnal geckos, with the former mostly preying on Formicidae, while the latter feeds on moths, small dipterans, and orthopterans (pers. obs.). We, therefore, predict that such potential prey will be more abundant near fire than under dark conditions, acting as the control.

## Materials and Methods

### Sampling and Study Area

The study was conducted in Southern Italy (40°15′N, 14°54′E, Salerno province) (Figure 2). The region is characterized by Mediterranean vegetation, predominantly featuring olive tree cultivation, rural structures, and stone walls.

**Figure 2.**
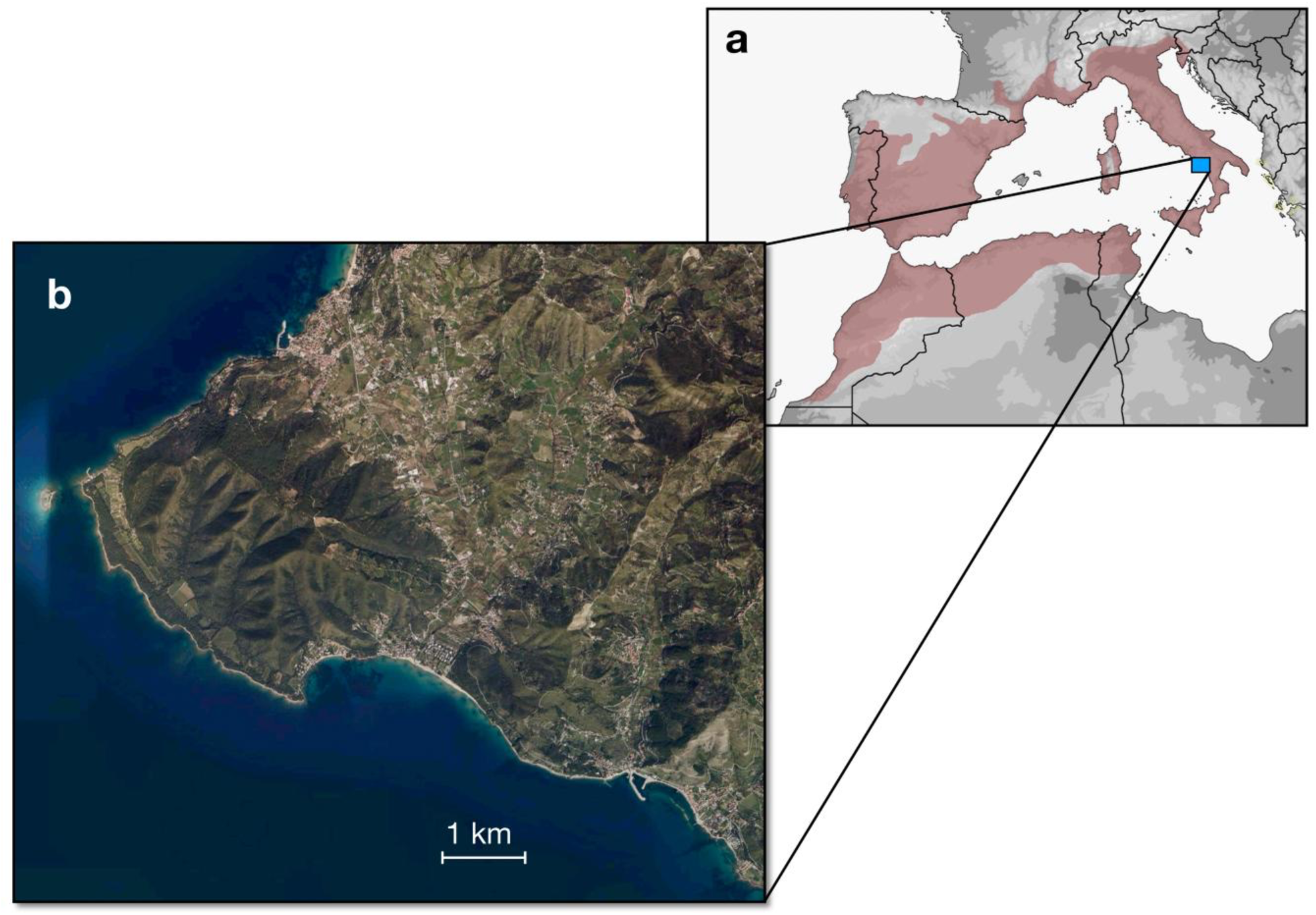
Location of the study area where the two forms of *Tarentola mauritanica* were sampled for the present study and the field experiment on the effect of fire on prey availability was set. The pink area in the general map shows the species’ current distribution.

Using nylon loops, we sampled 16 individuals: 6 dark-diurnal geckos collected from tree trunks and 10 pale-nocturnal geckos captured on walls. For each individual, we collected a section of the tail following autotomy and preserved it in a sterile tube containing 70% Et-OH, stored at -20°C until processing in the laboratory. Following tissue collection, the animals were released at the capture site, except for 6 individuals (3 dark-diurnal geckos and 3 pale-nocturnal geckos) sacrificed to obtain tissues suitable for histological analyses. We sampled Moorish geckos with the approval of the relevant nature conservation authority (Cilento, Vallo di Diano e Alburni National Park, prot. 2013/0010678). The experimental protocols were approved by the Ethical Committee for Animal Experiments at the University of Naples Federico II (protocol 2013/0032826).

### Skin Histology

We anaesthetised three dark-diurnal and three pale-nocturnal geckos with 250 mg/kg 1% MS-222 (Sigma Chemical Co. St. Louis, MO) injected into the intracoelomic cavity and euthanized them by decapitation. Following this, we processed skin samples taken from the backs of three dark and three light geckos for light microscopy, following the methods outlined in [34]. In brief, we fixed the samples in 2% paraformaldehyde and 2.5% glutaraldehyde (for 4 hours at 4°C) and subsequently post-fixed them in a 2% osmium tetroxide solution (for 1 hour at 4°C). After dehydration in an ascending series of ethyl alcohol, the samples were embedded in Epon 812 resin (Fluka). We then cut semi-thin sections of 1.5 μm thickness using a Super Nova Leica Ultratome and stained them with 1% toluidine blue in 1% sodium tetraborate buffer. Finally, we examined the sections using a Zeiss Axiocam camera attached to a Zeiss Axioskop microscope (Zeiss, Jena, Germany).

### Skin reflectance

To objectively determine the skin colouration of the sampled Moorish geckos, we measured skin reflectance using spectrophotometry (250–1000 nm, AvaSpec-2048-USB2-UA-50; Avantes, Apeldoorn, Netherlands), focusing on the range between 300 and 700 nm [25]. A white reference tile (WS2; Avantes) served as a calibration reference.

The spectrophotometer probe, featuring a 0.2 mm hole end, was positioned perpendicular to the animals’ body surface, and the reflectance (R%) was recorded at three locations on their backs. Subsequently, the average of the integrals subtending the reflectance curves was considered representative of the entire back of each individual [22,25,35]. We assessed the difference between the dark-diurnal and pale- nocturnal groups using a one-way ANOVA test, with statistical significance defined as p < 0.05.

### Melanocyte Stimulating Hormone (α-MSH) and Melanin Assay

Plasma and skin samples were obtained from three nocturnal/pale and three diurnal/dark geckos. Blood was drawn using a heparinized syringe from the interdiscal vertebra windows of the tail. Plasma was then isolated by centrifugation at 2000 g for 10 minutes at room temperature, and the samples were stored at −80 °C until processing in the laboratory.

We determined the levels of α-MSH using the ELISA assay as described by [36]. Statistical significance differences (p < 0.05) between the two groups were assessed using the Student’s t-test. To quantify melanin, we followed the protocol proposed by [37], with some modifications. Tissues were collected in 50 mM Tris-HCl pH 7.4, 300 mM NaCl, 0.5% NP-40, and protease inhibitors (Roche). Cells were lysed using a combination of freeze-thawing (3 cycles of dry ice at -37 °C), Dounce homogenization (200 strokes), and sonication (2 minutes, 10 seconds on and 10 seconds off).

### Phylogenetic Characterisation and Time Tree

To assess the haplotype diversity of sequenced mitochondrial genomes, we aligned them in Geneious version R9.16 with sequences of *T. mauritanica* (NCBI accession numbers from JQ300539 to JQ301443), representing distinct lineages [21], using *Gekko gecko* (accession number AY282753) as the outgroup.

To infer the divergence time of pale-nocturnal and dark-diurnal Moorish gecko populations, we constructed a time tree using MEGA11 (Tamura et al., 2021). We used sequences from 10 pale-nocturnal and 6 dark-diurnal geckos, along with a *T. mauritanica* sequence from Clade I (accession number JQ425050) as the outgroup. Additionally, we included a *T. mauritanica* Clade II sequence (accession number JQ425041) to enhance temporal resolution, as Clades I and II are temporally proximate [21].

Before constructing the time tree, we evaluated the haplotypes for JQ425041 and JQ425050 by aligning them with *T. mauritanica* sequences from JQ300539 to JQ301443 [21], representing diverse *T. mauritanica* lineages.

Initially, we performed an alignment with Muscle, considering the 18 mtDNA sequences. The following parameters were applied: gapopen = -400.00, gapextend = 0.00, max iterations = 16, using the UPGMA clustering method. To determine the most appropriate substitution model, we conducted a model selection test, opting for the Tamura-Nei method (BIC: 52037.98) based on the lowest Bayesian Information Criteria (BIC) score for constructing the phylogenetic tree.

A time tree was then inferred using the RelTime method [38,39] on the 18TN93Tree dataset in MEGA11 [40], comprising 16,618 positions. Branch lengths were calculated using the Maximum Likelihood (ML) method and the General Time Reversible substitution model [41].

The time tree was computed with one calibration constraint. Evolutionary rates from the ingroup were used to calculate divergence times, and times were not estimated for outgroup nodes, as the RelTime method does not assume that evolutionary rates in the ingroup clade apply to the outgroup. The estimated log-likelihood value of the tree is -25,781.91. A discrete Gamma distribution (5 categories; +G; parameter = 0.6794) was applied to model evolutionary rate differences among sites. To validate our findings from MEGA11, we also conducted divergence time estimation using BEAST v1.14 [42], with the following parameters: Substitution Model = TN93, Gamma Category Count = 5, Strict Clock Model, and setting priors to Coalescent Constant Population.

### Modelling potential human-gecko co-occurrence at the time the two gecko forms diverged

To establish whether Moorish geckos and humans coexisted when the dark-diurnal and pale-nocturnal geckos split, we generated Species Distribution Models (SDM) for *T. mauritanica* and projected them to the past. We collected modern occurrence data for the gecko from the Global Biodiversity Information Facility (GBIF) database by selecting “human observation” and “machine observation” as the basis of the record (www.GBIF.org, 26 January 2024; GBIF Occurrence Download https://doi.org/10.15468/dl.xprybx). The data were further filtered by excluding occurrences without geographical coordinates. For the human dataset, we used the *H. sapiens* records published by [43] and [44], focusing on fossil presence observations dated within a time window coherent with the divergence time estimates between the gecko lineages we found (see below). Radiocarbon data were calibrated using the “Bchron” R package [45] through the “intcal20” curve [46]. As environmental predictors, we employed the monthly bioclimatic variables generated through the 2Ma CESM1.2 simulation [44], downscaled at a 0.5° × 0.5° grid resolution. The native set of predictors was converted into bioclimatic variables according to WorldClim using the “dismo” R package [47]. Lastly, variables were projected onto the Mollweide coordinate reference system. To prevent model overfitting, duplicate occurrences in the raster grid were removed. After this step, we gathered 520 modern *T. mauritanica* and 740 past *H. sapiens* occurrences.

We defined the Mediterranean region as the study area for both *H. sapiens* and *T. mauritanica*, according to the current and historical geographical distribution of the Moorish gecko [21]). Then, we randomly generated 10,000 background points within the study area. To account for potential sampling biases, pseudoabsences were geographically placed according to the density of the occurrence data, making them more abundant where presences are denser [48–50]. For *H. sapiens* only, the record was divided into 1000-year consecutive time bins according to the time resolution of bioclimatic predictors. Subsequently, we partitioned the pseudoabsences proportionally to the number of presences per time bin. After extracting climatic values at each occurrence and pseudoabsence data point, we accounted for multicollinearity among predictors by considering the Variance Inflation Factor (VIF). The latter was assessed using the function “vifcor” embedded in the “usdm” R package, selecting a threshold of 0.75 [51,52]. After applying VIF, the selected predictors were BIO4 (temperature seasonality), BIO8 (mean temperature of the wettest quarter), BIO9 (mean temperature of the driest quarter), BIO13 (Precipitation of Wettest Month), BIO14 (precipitation of the driest month), BIO15 (Precipitation Seasonality), and BIO19 (precipitation of the coldest quarter).

To estimate the geographic distribution of both species in the past, we adopted the SDM ensemble approach. Specifically, we trained SDMs through an ensemble forecasting approach, as implemented in the R package “biomod2” [53]. We considered four different algorithms: Maximum Entropy Models (MaxEnt), Generalized Boosted Models (GBM), Random Forests (RF), and Generalized Linear Models (GLM). For model tuning, we adopted the default settings described in the “biomod2” R package. To evaluate the predictive accuracy of SDMs, we randomly split the dataset into a 70% sample used for model calibration and the remaining 30% used to evaluate model performance. Then, we calculated the area under the receiver operating characteristic curve (AUC; [54]) and the true skill statistic (TSS; [55]). This entire procedure was iterated 10 times, changing the randomly selected training/testing data points at each iteration. Model averaging was performed by weighting the individual model projections by their AUC values and averaging the results [56], excluding the model with AUC < 0.7. Lastly, SDM predictions were projected into the past to obtain the potential distribution of *T. mauritanica* in the Mediterranean area during the time interval predicted by the phylogenetic reconstruction as the divergence time between the two Moorish gecko forms.

To test whether there was a significant spatial correlation between *H. sapiens* and *T. mauritanica*, we adopted the Boyce Index [57]. It requires presence data points only and measures how much model predictions differ from a random distribution of the observed presences across the prediction gradients [58]. The Boyce Index varies between -1 and +1, whereby positive values indicate a model whose present predictions are consistent with the distribution of presences in the evaluation dataset, values close to zero mean that the model is not different from a random model, and negative values indicate counter predictions, i.e., predicting poor-quality areas where presences are more frequent [57].

### Testing differences in prey availability caused by fire

Within the study area, we selected two light-grey walls measuring 4x4 square meters each. One wall featured a fire at its base, while the other remained without a fire and was used as the control. Throughout 5 hours at night, we manually captured and identified all Arthropoda resting on the two walls. Identification involved examining diagnostic morphological traits using a Leica EZ4 W stereomicroscope and referencing a specialized manual [59] and local reference collections. We recorded the Orders and corresponding number of individuals, and then determined diversity and richness using Past3 software [60].

## Results

### Histological skin analysis

The comparison of histological skin samples taken from the dorsal surface of pale-nocturnal and dark- diurnal Moorish geckos revealed significant differences, in agreement with our hypothesis. Dark phenotype geckos exhibited numerous, predominantly large melanophores. Additionally, a thick and continuous layer of pigment granules was observed in the connective tissue of the underlying dermis. In contrast, the skin of pale-nocturnal specimens displayed remarkably reduced melanophores in size and abundance, and the layer of pigment surrounding them was sparse (Figure 3).

**Figure 3.**
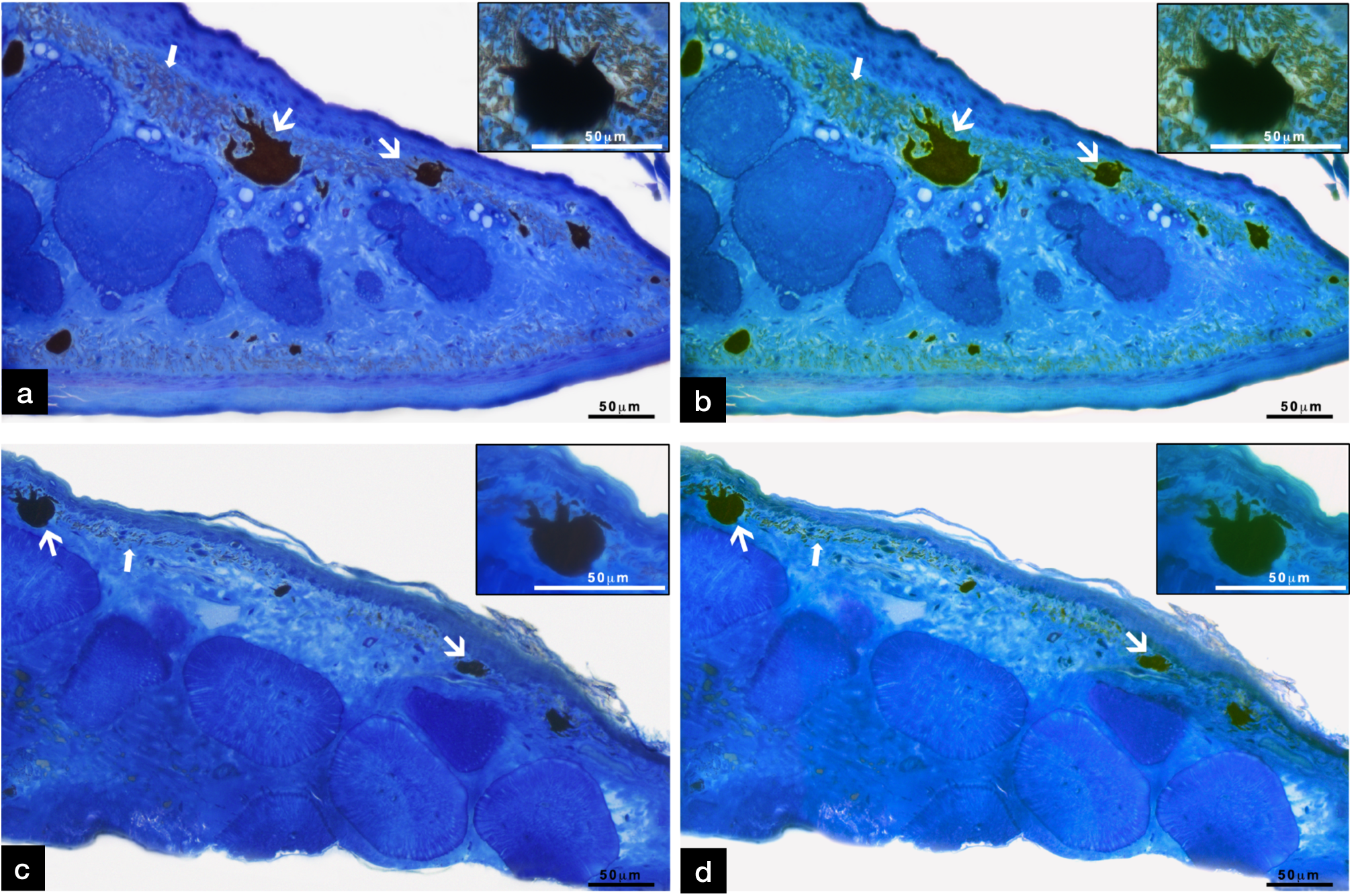
Skin sections stained with toluidine blue. a) Skin from the dorsal area of dark-diurnal Moorish geckos (*Tarentola mauritanica*), with single melanophores (inset). A high concentration of melanophores (thick arrow) is localized in the dermal region, along with a thick layer of melanin granules (arrow). b) Digitally stained version of image A to emphasise melanophores and melanin pigment. c) Skin from the dorsal area of a pale-nocturnal gecko, with a single melanophore (inset). d) Digitally stained version of image C to emphasise melanophores and a layer of melanin pigment.

### α-MSH and Melanin assay

Blood levels of α-MSH were significantly lower in pale-nocturnal geckos than in the dark-diurnal form (Figure 4a), resulting in a significant reduction in melanin production in the dermis of pale-nocturnal geckos (Figure 4b). This decrease in melanin production, consequently, led to a significantly smaller skin’s reflectance of pale-nocturnal geckos (Figure 4c).

**Figure 4.**
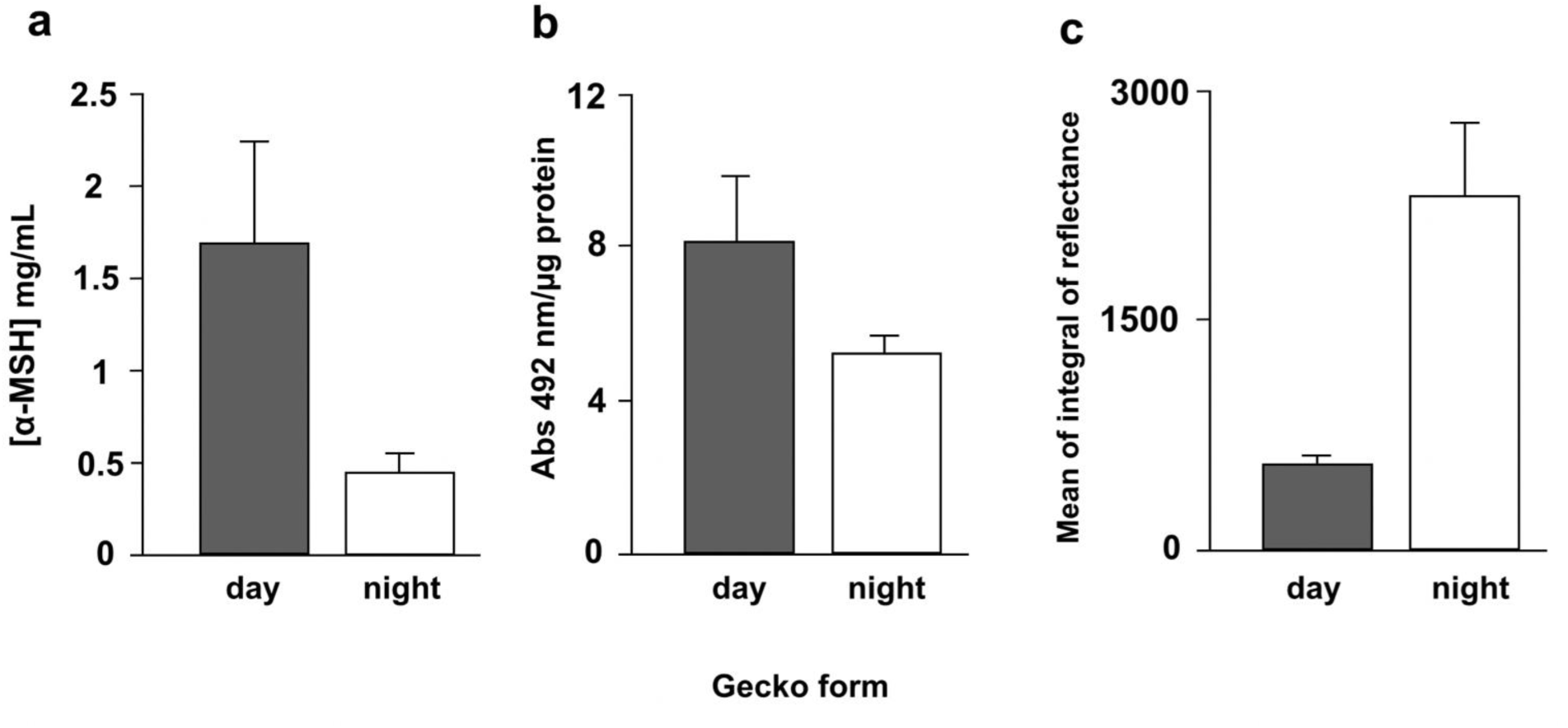
Comparisons of a) α-MSH hormone blood concentration, b) skin melanin assay, and c) skin reflectance between dark-diurnal (“day”) and pale-nocturnal (“night”) *Tarentola mauritanica* (one-way ANOVA tests, p<0.05, d.f. = 19, for all comparisons). Error bars represent the standard deviation.

### Phylogenetic reconstruction and molecular dating

The phylogenetic analysis of the Moorish gecko populations, adopting the complete mitochondrial genome, revealed their monophyletic nature, with the common ancestor estimated to have emerged approximately 500 thousand years ago (0.28-0.71 Mya) (Figure 5). Furthermore, the topology demonstrated a clear dichotomous division into the two monophyletic clades dark-diurnal and pale- nocturnal. The pale-nocturnal lineage was older (0.04 Mya) than the dark-diurnal (0.02 Mya) lineage.

**Figure 5.**
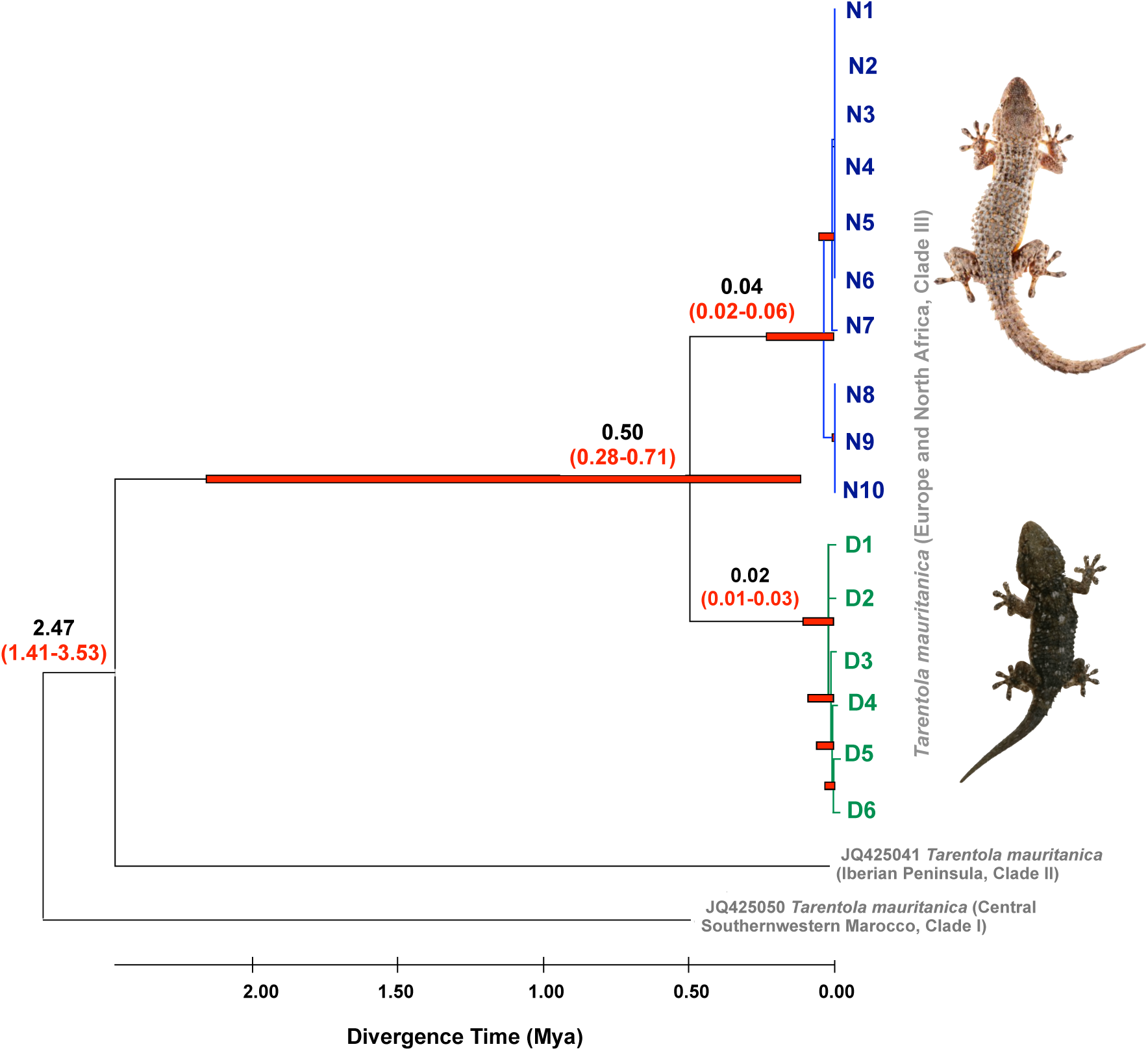
Phylogeny and divergence time estimation derived from molecular-clock analysis of 10 pale- nocturnal (N1-N10) and 6 dark-diurnal (D1-D6) *Tarentola mauritanica*. Divergence times for the two gecko populations were calculated using the complete mitochondrial genome. The red bars on the nodes represent the 95% credibility intervals of the estimated posterior distributions of the divergence times. Mya, million years ago.

### Modelling potential human-gecko co-occurrence at the time the two gecko forms diverged

The SDMs developed for humans and geckos were projected into the past to estimate their potential distribution in the Mediterranean area during the 60-20 kyr interval including the estimated time of divergence between the dark and pale gecko forms. These models exhibited high predictive performance. Specifically, the SDM developed for humans showed an AUC of 0.818 (S.D. = 0.014) and a TSS of 0.502 (S.D. = 0.028). Similarly, the SDMs developed for *T. mauritanica* achieved high performance, with an AUC of 0.96 (S.D. = 0.002) and a TSS of 0.803 (S.D. = 0.008). We assessed the spatial correlation between humans and the Moorish gecko in the past using the Boyce Index, obtaining a Spearman correlation value of 0.422. Furthermore, the linear mixed-effects model revealed a positive and significant (p < 0.001) relationship between *H. sapiens* presence and *T. mauritanica* potential distribution (Figure 6). Additionally, the model exhibited a relevant age-driven random effect structure (P = 0.016).

**Figure 6.**
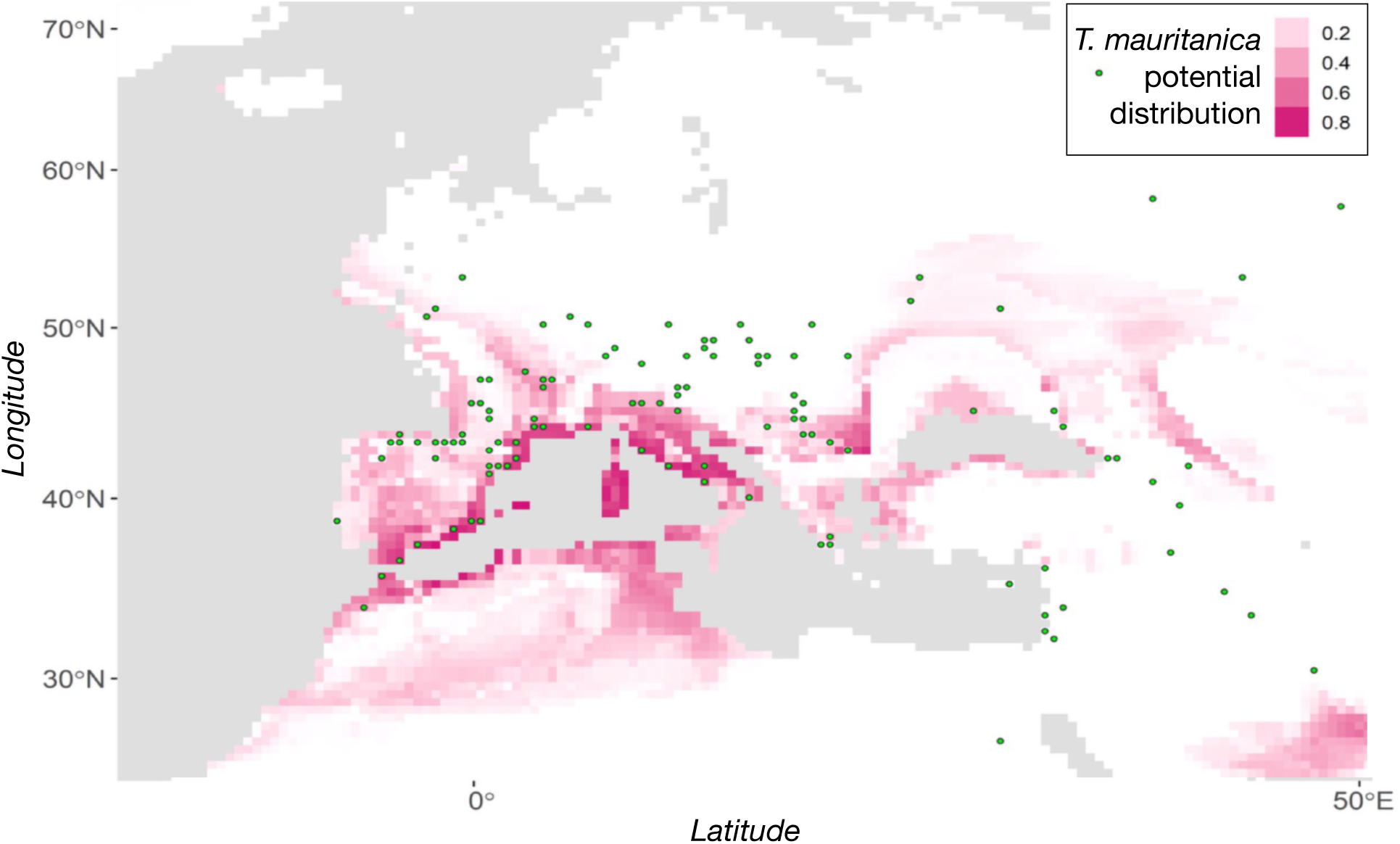
Spatial distribution model of *Tarentola mauritanica* and *Homo sapiens* during the time interval 60- 20 ky ago. The model indicates significant overlap in potential distribution within the Mediterranean region, supported by a linear mixed-effect model demonstrating a positive relationship between *H. sapiens* (depicted by green dots) and *T. mauritanica* potential distribution (highlighted in red).

### Testing differences in prey availability caused by fire

The species richness and abundance (number of total individuals) of arthropods sampled at the wall illuminated by fire differed significantly from those recorded at the unlit wall (Figure 7). Specifically, higher values were recorded at the lit wall for both arthropod richness (χ² = 6.250, d.f. = 1, *p* = 0.0124) and abundance (χ² = 38.111, d.f. = 1, *p* < 0.0001). The arthropod sample was dominated by moths, followed by orthopterans and spiders, whereas the less diverse and numerically inferior arthropod sample collected at the control wall only featured dipterans, spiders and ephemeropterans.

**Figure 7.**
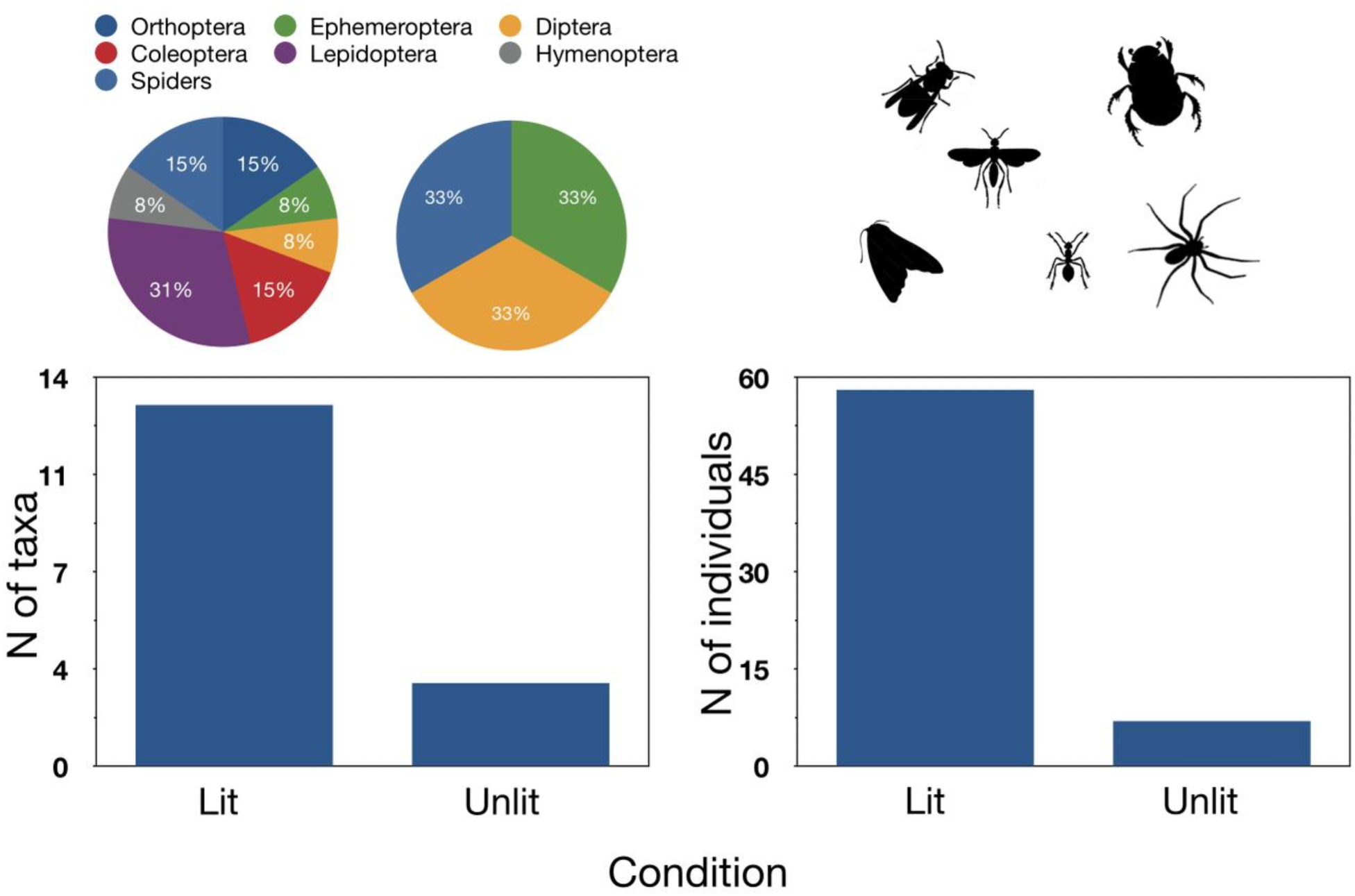
Differences in species richness (N taxa) and abundance (N individuals) of arthropods sampled at the illuminated wall compared to the unlit wall. Higher values were recorded at the lit wall for both arthropod richness (*p* = 0.0124) and abundance (*p* < 0.0001).

## Discussion

### May human-controlled fire have influenced the temporal niche partitioning in Moorish geckos?

We used a multidisciplinary approach to demonstrate that the two gecko variants have fixed their phenotypes over the evolutionary path and that the nocturnal-pale form has smaller and less numerous melanosomes and is subject to lower α-MSH levels compared to the diurnal-dark form. Moreover, using a full mitochondrial genomic analysis, we found that the pale-nocturnal lineage evolved 20,000 years before the dark-diurnal lineage and that both forms appeared at a time when modern humans invaded Europe bringing with them widespread control of fire, which must have represented a key attractor of suitable arthropod food for the pale geckos, thus providing a novel, previously unexploited foraging niche. Our modelling exercise confirmed sympatry between humans and geckos when the two lineages arose. Finally, a simple experiment showed that fire lit near a clear wall increases considerably the amount of arthropod food, attracting selectively insects such as moths and orthopterans that are elective prey for the pale variant, unlike those eaten by the dark form in daytime foraging.

Overall, the multifaceted body of evidence provided suggests that ancient fires used by humans may have facilitated the separation between the two gecko variants leading to the situation we observe today, yet it does not fully clarify whether the two lineages originated from diurnal or nocturnal ancestors. While human-controlled fire may have represented an environmental novelty providing an important foraging niche at night, the evidence we gathered is still compatible with either scenario. Our phylogenetic tree marks the appearance of *T. mauritanica* 50,000 years ago (range 28,000-51,000 years ago), which is compatible with the presence in Italy of this species during the Upper Pleistocene supported by fossil remains [61] and rules out a historical, anthropogenic introduction of the species in the country, in agreement with previous phylogenetic work [62]. The timing of human control over fire remains a contentious issue [63,64]. Evidence from Europe suggests that Neanderthals and early modern humans used controlled and consistent fire starting from the Middle Pleistocene onwards in Europe, and from the Late Pleistocene in Europe and South Africa, for purposes beyond warmth and cooking [65,66]. However, the hypothesis that controlled fires became prevalent upon the arrival of modern humans in Europe, particularly in the Mediterranean, is highly plausible.

### The ”out of the dark” scenario

The hypothesis of a nocturnal ancestor that has led to a first nocturnal lineage and only 20,000 years later to a diurnal descendant is certainly parsimonious and agrees with the fact that based on a phylogenetic analysis of temporal activity patterns, nocturnality would be an ancient trait appearing at the base of the gecko tree [67].

Geckos and skinks independently evolved nocturnality, diverging from the ancestral diurnality typical in lizards [68], with noticeable metabolic [69,70] and sensory [71] adaptations to the nocturnal niche. While the spread of human-controlled fire may have represented a strong advantage for more night-specialized phenotypes, these may have monopolized such preferred foraging sites excluding other less competitive individuals. In behavioural studies of *T. mauritanica* foraging near artificial lights, large individuals exclude smaller and younger geckos from key foraging sites [72], and a similar situation is conceivable near the fires set by ancient humans. On the other hand, leaving on the periphery of such sites for subordinate geckos constrained in their marginal territories may have been highly unprofitable because of the “vacuum effect” caused by the fire, attracting arthropod food and depleting the surrounding areas. This situation is well-known today in response to artificial lighting and is seen as one of the adverse effects of light pollution on natural environments, favouring a few opportunistic, light-tolerant predators at the expense of light-averse species whose dark habitat is continuously impoverished [10,73]. The process has been proposed as the way light-tolerant common pipistrelles (*Pipistrellus pipistrellus*) have outcompeted light-averse lesser horseshoe bats (*Rhinolophus hipposideros*) in Switzerland, contributing to the decline of the latter species [74]. Under such a scenario, the most phenotypically plastic individual geckos capable of acquiring a sufficiently dark colour to shift their temporal niche to the daytime might have led to the evolution of the diurnal lineage observed today, escaping the strong competition by the nocturnal dominants.

### The ”into the dark” scenario

The alternative “into the dark” scenario we considered would originate from a dark-diurnal ancestor from where the pale-nocturnal lineage, and, 20,000 years later, the dark-diurnal lineage would evolve. During the evolutionary path leading to the current nocturnal specialists, an ancestral dark form lost the ability to change conspicuously its coloration, fixing a pale phenotype that would effectively mitigate predatory pressure at night [22]. The attainment of the pale phenotype involved a reduction in α-MSH hormone levels, as well as alterations in the number and size of melanosomes in the dermis. In other words, predators would have exerted strong selective pressure, killing all individuals whose colouration was not pale enough to fade their contour on rock walls at night, leaving only the palest ones on the scene. The paleo-synanthropic relationship that took place between geckos and humans would have been carried forward until modern times when the nocturnal-pale gecko inhabits built-up areas and often sits and waits for its prey near artificial sources of lighting.

Although nocturnal lizards such as geckos may be active at body temperatures that are considerably lower than those characterizing activity in diurnal lizards [75,76], nocturnal species may still experience suboptimal locomotion in the cold of the night [75–77], which might, in theory, limit successful foraging. However, locomotion in nocturnal lizards is surprisingly efficient, as they outrun threefold diurnal lizards and show low-temperature performances like those of diurnal lizards at higher temperatures, thanks to a low minimum cost of locomotion [75,78] and higher metabolic rates at low temperatures (Hare et al., 2010). In the case of pale geckos, hunting near fires might have represented a further way of warming up and achieving even higher locomotion performances.

The role of phenotypic plasticity in promoting the diversification of new lineages has been reconsidered in recent times [79], and today is seen as potentially important, yet its mechanisms are still debated (e.g., [36,80–84]. Phenotypic plasticity might serve as a protective mechanism against environmental fluctuations, fostering the persistence of populations [79,85,86]. On the other hand, phenotypic plasticity could lay the groundwork for a process called “genetic assimilation,” where a phenotype becomes integrated into the genotype, potentially resulting in a loss of plasticity known as “canalization” [87]. Under the “into the dark” scenario, the plasticity still observed in the diurnal-dark lineage and hypothetically shared with its diurnal ancestor facilitated the initial colonization and survival of the population in a new environment, allowing time for subsequent genetic adaptation to fine-tune responses to this environment. The plastic capabilities of the phenotype can thus respond to selection, anticipating the adaptation process of the genotype and enabling the colonization of the new nocturnal niche.

Current artificial lighting at night is offering numerous tests of how animals respond to novel sources of light [12], and this also holds for geckos. For instance, six gecko species from the genus *Phelsuma*, although mostly diurnal, shifted their activity from diurnal to nocturnal hours to exploit prey concentrating near artificial lights [88], a behavioural change closely resembling that we propose as a potential origin of the Moorish gecko’s nocturnal lineage.

### The influence of human-controlled fire on temporal niche partitioning in Moorish geckos

In conclusion, our investigation into the potential influence of human-controlled fire on the temporal niche partitioning in Moorish geckos presents intriguing insights but also leaves some questions unanswered. The multifaceted evidence we gathered suggests that ancient fires used by humans may have played a role in facilitating the separation between the two gecko variants observed today. However, it does not definitively clarify whether the two lineages originated from diurnal or nocturnal ancestors. The “out of the dark” scenario posits a parsimonious hypothesis where a nocturnal ancestor led to a first nocturnal lineage, followed later by a diurnal descendant, whereas the “into the dark” scenario suggests an alternative origin, where a dark-diurnal ancestor gave rise to a less plastic, more specialised pale- nocturnal lineage. The role of phenotypic plasticity emerges as a crucial factor in both scenarios, potentially facilitating the colonization of new environments and the subsequent fine-tuning of genetic adaptations.

The ongoing impact of artificial lighting on nocturnal behaviour underscores the relevance of our findings, as it offers parallels to the adaptation of the Moorish gecko’s nocturnal lineage. Future research could explore further the mechanisms underlying phenotypic plasticity and its interplay with genetic adaptation in this species, shedding light on the evolutionary dynamics of temporal niche partitioning in response to past and current human influences.

## Author Contributions

Conceptualization: D.F. and M.B.; Sample collection: D.F., E.R., V.M., and M.B; Laboratory experiments: E.R., V.M., B.A. and M.B; Analyses of data: D.F., D.R., E.R., B.A., A.M., G.G., and M.B; Supervision: D.F.; Writing original draft: D.F., E.R. and M.B.; Review & editing: D.F., D.R., E.R., V.M., B.A., A.M., G.G. and M.B. All the authors read and approved the manuscript.

## Data Availability

The sequencing data are deposited in GenBank under accession numbers MK275668 - MK275687

## Acknowledgments

We thank the Professor Greger Larson for suggestion and revision of the manuscript.

## Graphical abstract

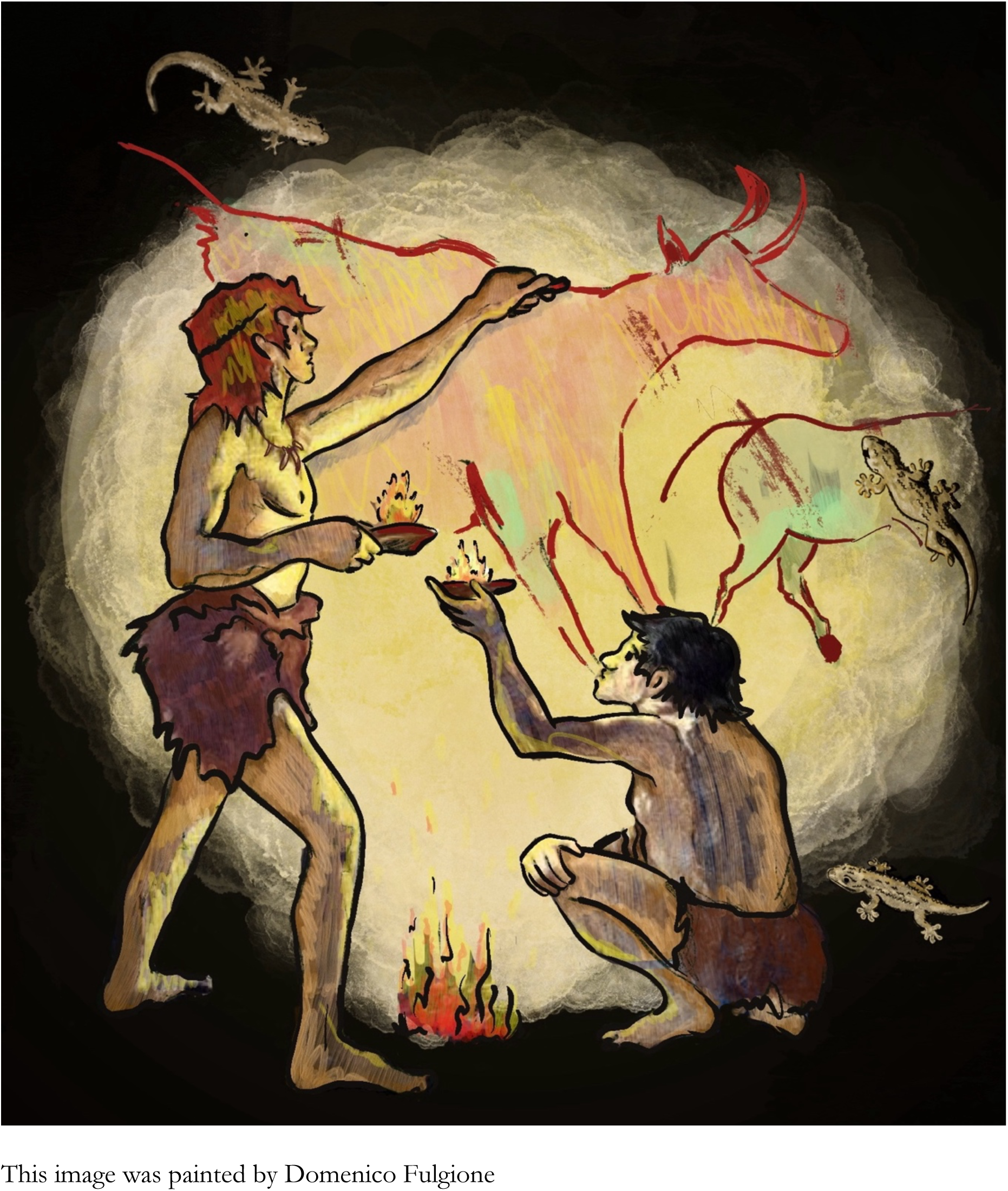

